# Large protein complex interfaces have evolved to promote cotranslational assembly

**DOI:** 10.1101/2021.05.26.445863

**Authors:** Mihaly Badonyi, Joseph A Marsh

## Abstract

Assembly pathways of protein complexes should be precise and efficient to minimise misfolding and unwanted interactions with other proteins in the cell. One way to achieve this efficiency is by seeding assembly pathways during translation via the cotranslational assembly of subunits. While recent evidence suggests that such cotranslational assembly is widespread, little is known about the properties of protein complexes associated with the phenomenon. Here, using a combination of proteome-specific protein complex structures and publicly available ribosome profiling data, we show that cotranslational assembly is particularly common between subunits that form large intermolecular interfaces. To test whether large interfaces have evolved to promote cotranslational assembly, as opposed to cotranslational assembly being a non-adaptive consequence of large interfaces, we compared the sizes of first and last translated interfaces of heteromers in bacterial, yeast, and human complexes. We detect a clear preference in N-terminal interfaces, which will be translated first and thus more likely to form cotranslationally, to be larger than C-terminal interfaces, suggesting that large interfaces have evolved as a means to maximise the chance of successful cotranslational subunit binding.

## Introduction

The majority of proteins across all domains of life function as part of multimeric complexes. Although we have a comprehensive understanding of the diverse quaternary structure space occupied by complexes (Ahnert *et al*., 2015), much less is known about where, when, and how their component subunits assemble. Continuing advances in cryo-electron microscopy, mass photometry, and genetic interaction mapping (Braberg *et al*., 2020) are facilitating a transition towards a structural view of proteomes (Levy and Vogel, 2021). While, at the present, our structural analyses are primarily limited to structures of complexes in the Protein Data Bank (Berman *et al*., 2000), there is a new generation of multiscale protein complex modelling approaches (Evans *et al*., 2021; Humphreys *et al*., 2021; Gao *et al*., 2022) promising to fill the gap and accelerate structure-based discovery. Our understanding of proteomes has also been dramatically improved by the development of ribosome profiling, which has provided us with quantitative measurements at the level of translation. Alterations of the technique revealed the cotranslational action of chaperones (Oh *et al*., 2011; Shiber *et al*., 2018; Stein, Kriel and Frydman, 2019), shed light on the role of collided ribosomes in proteostasis (Arpat *et al*., 2020; Han *et al*., 2020; Zhao *et al*., 2021), and supported the view of the ribosome as a signalling hub (D’Orazio and Green, 2021). To the present work, however, it is of outstanding relevance that ribosome profiling has laid down strong evidence that the assembly of protein complexes often starts on the ribosome (Duncan and Mata, 2011; Natan *et al*., 2017; Sepulveda *et al*., 2018; Shiber *et al*., 2018; Kamenova *et al*., 2019; Panasenko *et al*., 2019; Bertolini *et al*., 2021).

Two factors appear to be particularly important for cotranslational assembly: the proximity of nascent chains on adjacent (*cis*) or between juxtaposed (*trans*) ribosomes, and the localisation of interface residues towards the N-terminus of a protein, which allows more time for an interaction to occur during translation (Shieh *et al*., 2015; Natan *et al*., 2018; Kamenova *et al*., 2019). Recent findings demonstrated that homomers, formed from multiple copies of a single type of polypeptide chain, frequently assemble on the same transcript via the interaction of adjacent elongating ribosomes (Bertolini *et al*., 2021). This mechanism is highly effective because it takes advantage of the fact that homomeric subunits are identical; therefore nascent chains are essentially colocalised by the nature of their synthesis. Although homomers may benefit from polysome-driven assembly, it requires allocation of cellular resources to ensure at least two ribosomes are actively translating the same mRNA at any one time (Liu, Beyer and Aebersold, 2016). On the other hand, heteromers, products of different genes that physically interact, can only employ the *trans* mode of assembly in eukaryotes, providing mechanisms that colocalise their transcripts exist (Liu *et al*., 2016; Pizzinga *et al*., 2019; Wang *et al*., 2020; Chen and Mayr, 2022). In contrast to homomers, cotranslational assembly of heteromers may only require a single ribosome on each mRNA, which could allow lowly abundant regulatory proteins to cotranslationally assemble (Heyer and Moore, 2016; Biever *et al*., 2020). Alternate ribosome usage and translation-coupled assembly can explain how cells achieve efficient construction of complexes with uneven stoichiometry, accounting for a substantial fraction of heteromeric complexes (Marsh *et al*., 2015).

Despite growing evidence supporting the importance of cotranslational assembly, far less is known about the properties of the interfaces involved. It has been observed that cotranslationally binding subunits have a tendency to fall out of solution or become degraded by orphan subunit surveillance mechanisms in the absence of their partner subunits (Choe *et al*., 2016; Juszkiewicz and Hegde, 2018; Natan *et al*., 2018; Shiber *et al*., 2018; Kamenova *et al*., 2019). This observation may be explained under two assumptions: N-terminal interfaces are aggregation prone due to interference with cotranslational folding (Ciryam *et al*., 2013; Jacobs and Shakhnovich, 2017; Kudva *et al*., 2018; Kramer, Shiber and Bukau, 2019), and/or that cotranslationally forming interfaces possess unique structural properties that predispose them to aggregation in the absence of binding partners. Whilst there is evidence for the former (Natan *et al*., 2018), interfaces involved in nascent chain assembly have not been systematically studied before. Therefore, we cannot exclude the possibility that they have structural features that make them more susceptible to a cotranslational route.

Hydrophobic surfaces play a key role in nucleation theory (Hermann, 1972; Tanford, 1978; Chandler, 2005) and protein folding (Privalov and Khechinashvili, 1974; Chothia, 1975; Gething and Sambrook, 1992), but more importantly, hydrophobicity remains the founding principle of protein-protein recognition theory (Kauzmann, 1959; Chothia and Janin, 1975). Defined as the buried surface area between subunits, interface size shows correspondence to hydrophobic area because larger interfaces contain more interface core residues (Levy, 2010). Conveniently, interface area is relatively simple to compute from structural data (Hubbard SJ, 1993; Kleinjung and Fraternali, 2005; Winn *et al*., 2011; Mitternacht, 2016). Whilst the relationship between interface area and measured affinity is non-linear (Eisenberg and Mclachlan, 1986; Horton and Lewis, 1992; Brooijmans, Sharp and Kuntz, 2002; Vangone and Bonvin, 2015), interface area shows remarkable correspondence with subunit dissociation energy, and is reflective of the evolutionary history of subunits within complexes (Levy *et al*., 2008).

We hypothesised that cotranslational interactions may be distinguished from others based upon the areas of the interfaces involved. The size hierarchy of interfaces in protein complexes can be used to predict the order in which their subunits assemble, in good agreement with experimental data (Levy *et al*., 2008; Marsh *et al*., 2013; Wells, Bergendahl and Marsh, 2016). According to this theory, the largest interfaces in a complex correspond to the earliest forming subcomplexes within the assembly pathway, irrespective of the binding mode. While specific contacts that increase affinity would introduce compositional biases into the sequence space, exerting undue selection pressure on proteomes, variability in interface size can emerge from non-adaptive processes as the organising principle of cotranslational assembly (Conant, 2009; Ahnert *et al*., 2010; Gray *et al*., 2010; Lynch, 2013; Leonard and Ahnert, 2019; Hochberg *et al*., 2020).

In the present study, we address this idea by analysing experimental data on cotranslationally assembling human proteins (Bertolini *et al*., 2021). Our results establish a strong correspondence between cotranslational assembly and subunit interface size. To test whether large interfaces could represent an evolutionary adaptation to cotranslational assembly, we took advantage of the many protein complex subunits that have more than one interface. We compared the areas of first and last translated interfaces in bacterial, yeast, and human heteromeric subunits, and found a clear tendency for the first interface to be larger than the last interface across all species. This finding suggests that large protein complex interfaces have evolved to promote cotranslational assembly.

## Results

### Cotranslationally assembling subunits are characterised by large interfaces

In a recent study, a novel ribosome profiling method was used to identify over 4000 cotranslationally assembling human proteins (Bertolini *et al*., 2021). By design, the method can identify subunits that undergo cotranslational assembly when both subunits are in the process of translation. As recently proposed (Kamenova *et al*., 2019), we refer to this mode of binding as “simultaneous” assembly (**Figure 1A/B**).

**Figure 1.**
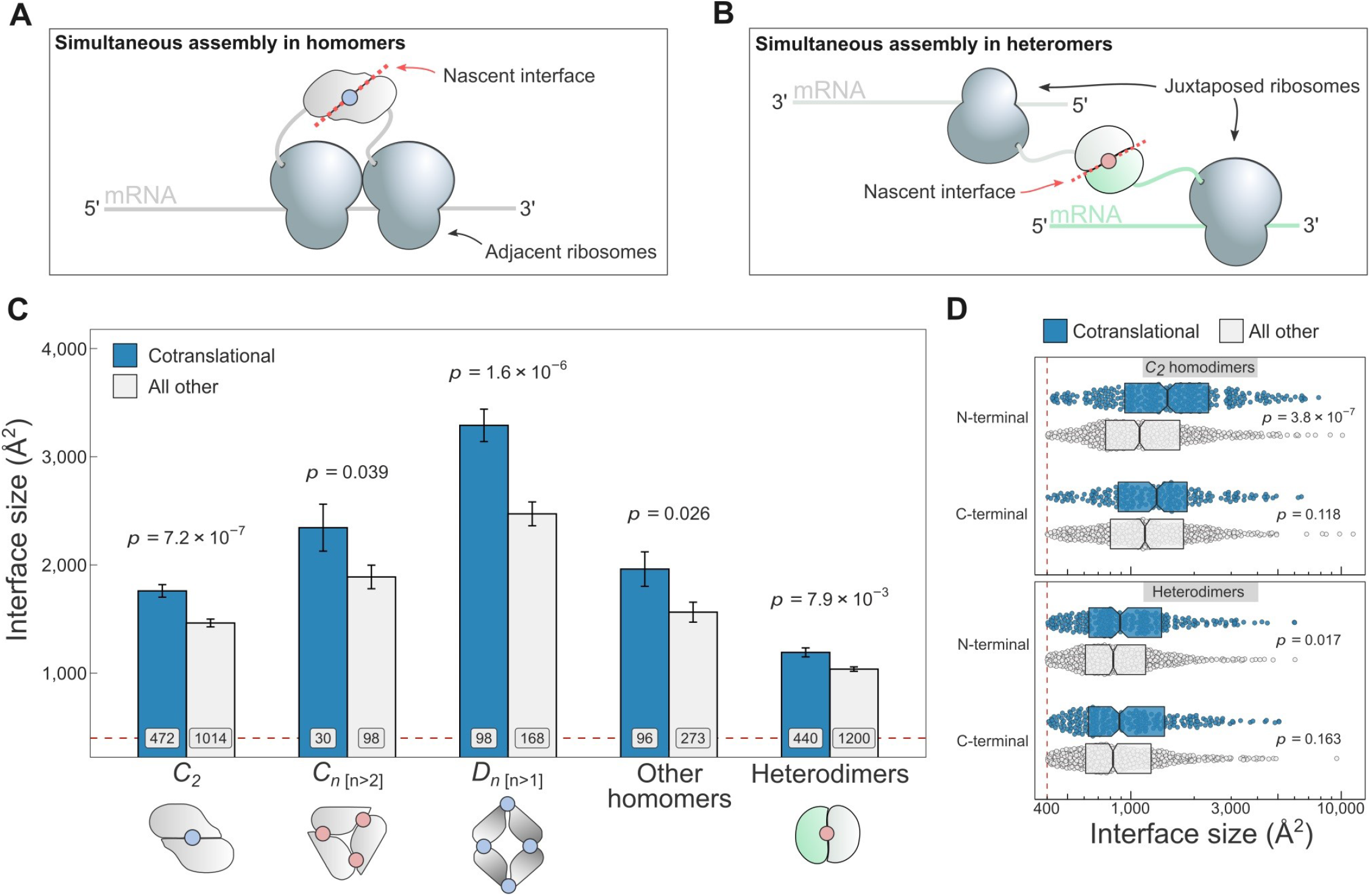
(**A**) Schematic representation of (*cis*) simultaneous cotranslational assembly in homomers. (**B**) Schematic representation of (*trans*) simultaneous cotranslational assembly in heteromers. (**C**) Interface size differences between cotranslationally assembling and all other subunits of homomeric symmetry groups and heterodimers. Error bars represent SEM and labels on bars show the number of proteins in each group. The *p*-values were calculated with Wilcoxon rank-sum tests. Pictograms show the basic structure of symmetry group members, with the blue dots representing isologous and red dots heterologous and heteromeric interfaces. (**D**) Interface size distributions of cotranslationally assembling and all other subunits of *C*_2_ homodimers and heterodimers, subset by the terminal location of the interface. The *p*-values were calculated with Wilcoxon rank-sum tests.

To investigate if interface area correlates with simultaneous assembly, we computed the buried surface areas of homomeric and heterodimeric subunits, and subset the results by whether or not the protein was detected to cotranslationally assemble. The arrangement of homomeric subunits with respect to one or more rotational axes allows their classification into symmetry groups. The three most common groups are the twofold symmetric (*C*_2_), cyclic (*C*_n [n>2_]), and dihedral (*D*_n[n>1_]) complexes, which all have distinct structural and functional characteristics (Goodsell and Olson, 2000; Levy and Teichmann, 2013; Bergendahl and Marsh, 2017) and should therefore be considered separately. In **Figure 1C**, we show the results of the interface size analysis broken down by the symmetry of the complexes.

*C_2_* homodimers represent the most highly populated symmetry group and their single isologous interface (*i.e*. symmetric or head-to-head) makes the analysis simple to perform. In line with our expectation, *C_2_* symmetric subunits that assemble during translation expose 20% larger areas than those that do not (*p* = 7.2 × 10^-7^, Wilcoxon rank-sum test). Considering that the cotranslational assembly annotations are derived from a laboratory technique that uses extensive biochemical fractionation, it is important to control for the possibility that larger interfaces would be more persistent to these procedures. We therefore controlled for the potential confounding effect of larger interfaces by setting incrementally higher interface area cutoffs (**Figure S1A**), to which the trend appears robust.

Higher-order cyclic complexes are centred on a rotational axis so that every subunit has two distinct interfaces, each with an adjacent protomer. Both interfaces are heterologous (*i.e*. asymmetric or head-to-tail) and approximately the same size. Cyclic symmetry is potentially confounded by its tendency to form ring-like structures, which are ubiquitous components of biological membranes (Forrest, 2015). As a result, membrane-bound complexes are enriched in non-polar amino acids that form the interface with the alkane core of the lipid bilayer. We focused on the analysis of cyclic homomers that do not localise to the plasma membrane, owing to competing hydrophobic forces exerted by protein-lipid interactions. Despite the limited number of structures available, we detect a significant difference in interface area among soluble members of the cyclic symmetry group (**Figure 1C**), with the mean of cotranslationally forming subunits being 24% larger (*p* = 0.039, Wilcoxon rank-sum test). Notably, we did not observe a trend in plasma membrane localised cyclic complexes (**Figure S1D**).

Dihedral symmetry can be thought of as the stacking of a dimeric or cyclic complex through the acquisition of a twofold axis. All dihedral complexes have isologous interfaces, and those with at least six subunits can have both isologous and heterologous interfaces (*e.g. D*_3_ dimers of cyclic trimers). We find that cotranslationally assembling dihedral complexes have on average 33% larger interfaces than those assumed to assemble after their complete synthesis (**Figure 1C**; *p* = 1.6 × 10^-6^, Wilcoxon rank-sum test). A dihedral complex is likely to have evolved from a *C*_2_ homodimer if its largest interface is isologous and, conversely, when its heterologous interface is largest, the complex probably arose via a cyclic intermediate (Levy *et al*., 2008; Marsh and Teichmann, 2014). When dihedral complexes are grouped by their likely evolutionary history, the trend is present in both groups (**Figure S1D**), consistent with that observed in *C*_2_ homodimers and higher-order cyclic complexes.

We pooled all remaining homomers, including those with helical and cubic symmetry, and those that are asymmetric, into a single “other” category, due to their relatively low representation in the human proteome. Altogether, cotranslationally assembling subunits in this heterogeneous category present 25% larger interface areas than other members (**Figure 1C**; *p* = 0.026, Wilcoxon rank-sum test). Thus, the interface size trend in cotranslationally assembling complexes appears to hold up across all types of homomers.

Because simultaneous assembly in heteromers requires two different transcripts positioned in *trans* (**Figure 1B**), we were curious if they, too, showed a correspondence between cotranslational assembly and interface size. Due to the diverse quaternary structures and assembly pathways associated with heteromeric complexes (Ahnert *et al*., 2015), we focused on the simplest cases, the heterodimers, which form a single heteromeric interface by the physical interaction of two different proteins. When compared, heterodimers that simultaneously assemble reveal a 15% larger interface area on average than those lacking annotations (**Figure 1C**; *p* = 7.9 × 10^-3^, Wilcoxon rank-sum test). Similar to *C*_2_ homodimers, the trend in heterodimers is also robust to incremental interface area cutoffs (**Figure S1A**), making it unlikely to be an experimental artefact.

The weaker effect size in heterodimers relative to *C_2_* homodimers may be explained by the combination of two factors. First, previous experimental evidence suggests that heteromers commonly employ the “sequential” mode of assembly, whereby a subunit in the process of translation recruits a fully synthesised and folded subunit (Shiber *et al*., 2018; Kamenova *et al*., 2019). This mode of assembly has not yet been experimentally probed on a proteome-wide scale, and it is possible that many heterodimers lacking cotranslational assembly annotations in our data set employ sequential assembly. As a result, assuming that interface size plays a role in sequential assembly as well, these unannotated proteins weaken the effect we can detect. Second, it is plausible that another biological process, yet uncharacterised in detail, is responsible for the colocalisation of transcripts and the subsequent subunit assembly (Wang *et al*., 2020; Chen and Mayr, 2022), which could make assembly in heteromers less reliant on interface area.

We considered three potentially confounding variables of the ribosome profiling method, which was used to detect cotranslationally assembling proteins. First is protein length, in part because long polypeptide chains take more time to translate, making it more likely for a cotranslational interaction to come about, and partly because bigger proteins tend to form larger interfaces. Long proteins are also encoded by long transcripts on which structures called di-ribosomes (two ribosomes connected by interacting nascent chains) may persist for a longer time period, potentially leading to their survivorship bias to the observer. In **Figure S1B**, we present an analysis where both *C_2_* homodimers and heterodimers are binned by their length into bins containing equal number of structures, and subset by cotranslational assembly. With the exception of long heterodimers, all bins follow the expected interface size trend. More importantly, for both types of complexes, the middle bin, which contains approximately 350-720 residue long proteins and thus covers a large fraction of the human proteome, shows the strongest effect size.

The second variable we accounted for is the confidence-based classification of the cotranslational assembly data set. Bertolini and co-workers employed an elaborate strategy to assign high or low confidence to the candidates (details in Bertolini et al., 2021). High confidence proteins only make up a fifth of all annotations, which prohibits their exclusive use in our analyses. However, in all homomeric symmetry groups and in heterodimers, both high and low confidence candidates have a larger mean area than unannotated proteins (**Figure S1C**).

The third potential confounder is the location of the interface relative to protein termini. Interactions via N-terminal interfaces are translated earlier, therefore increasing the time available for them to assemble cotranslationally. Given that cotranslationally forming interfaces identified by ribosome profiling are known to be significantly enriched towards the N-terminus of proteins (Bertolini *et al*., 2021), but that overall, homomeric interfaces tend to be enriched towards the C-terminus (Natan *et al*., 2018), we wished to control for interface location. We classified all interfaces as occurring on either the N- or C-terminal halves of proteins, based on the position of the interface midpoint, which is the residue at which half of the buried surface area of an interface is reached. This comparison is presented for *C_2_* homodimers and heterodimers in **Figure 1D**. In all groups, there is a clear interface size trend wherein cotranslationally assembling subunits have a larger area. More interestingly, however, the trend is only significant and much larger in effect between N-terminally localised interfaces. In fact, N-terminal interfaces are significantly larger than C-terminal interfaces in cotranslationally assembling homodimers (*p* = 0.021, Wilcoxon rank-sum test). One possible explanation for this is that N-terminally localised interfaces are far more likely to represent cases of genuine cotranslational assembly.

### Larger and earlier-assembling interfaces tend to form cotranslationally in heteromeric subunits with multiple interfaces

Having confirmed that subunit interface size correlates with cotranslational assembly, we next wanted to see if this trend applies within single subunits that have more than one interface. In other words, do multi-interface heteromeric subunits also employ their largest interface during the course of simultaneous assembly? A multi-interface heteromeric subunit forms at least two distinct interfaces with two other proteins in a complex that contains at least three subunits. Because of the interface hierarchy that exists within protein complexes (Levy *et al*., 2008; Marsh *et al*., 2013), we hypothesised that the largest interface, which is most likely to assemble earliest, should also be more likely to cotranslationally assemble.

To perform an analysis at the multi-interface level, we made use of the assembly-onset positions determined for every protein in the cotranslational assembly data set (Bertolini *et al*., 2021). An assembly-onset is a single residue in the protein sequence, whose codon is being decoded by the ribosome at the time of cotranslational assembly. In order to identify which interface the assembly-onsets belongs to, we mapped them to the closest interface midpoint in the linear protein sequence, as illustrated in **Figure 2A**. This is to avoid biases from large interfaces, which have many more interface residues and therefore a higher probability that an assembly-onset would map to them if the interface was not compressed into a single residue, the midpoint.

**Figure 2.**
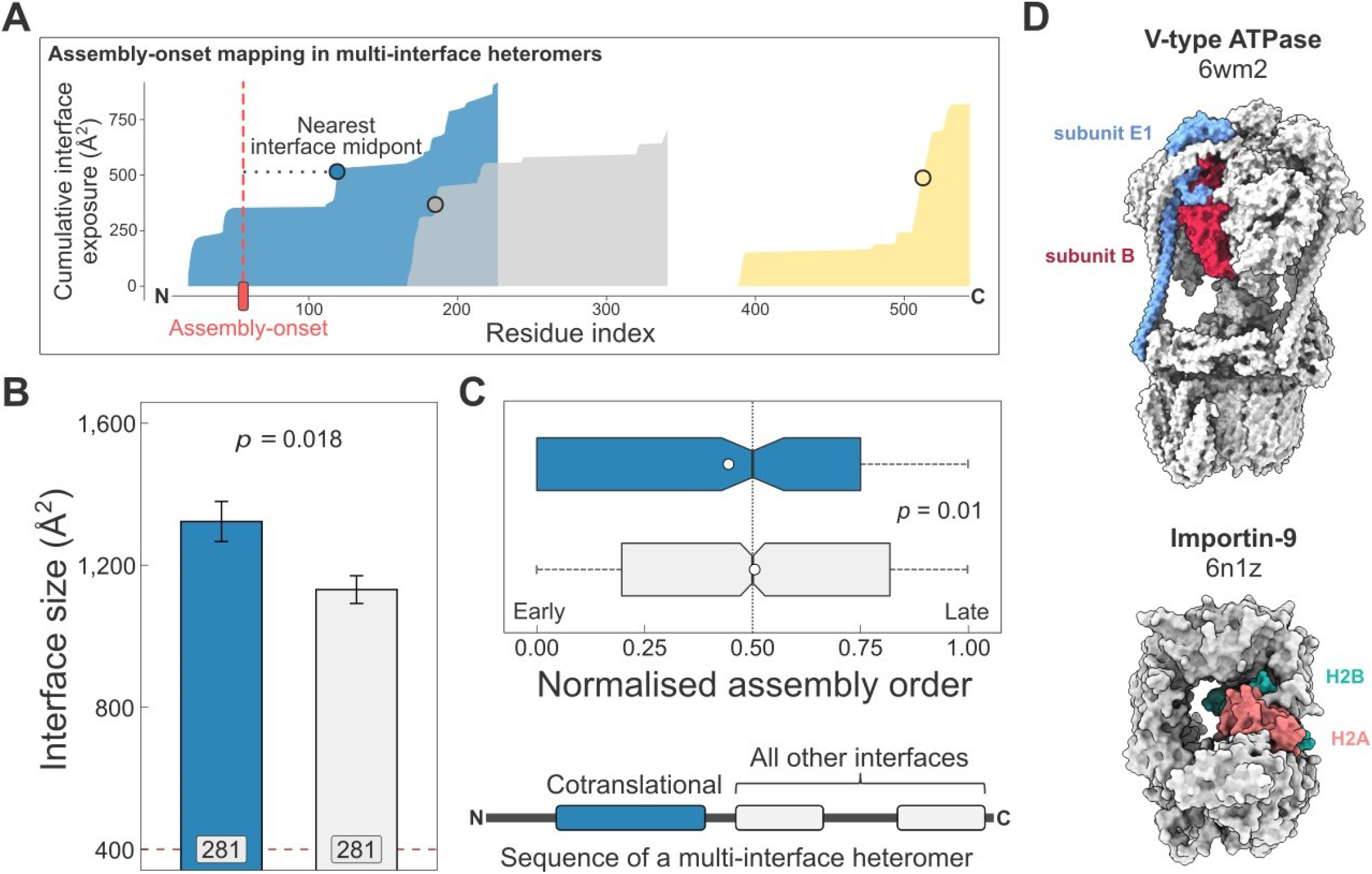
(**A**) Visual representation of the interface mapping protocol. Area charts show the cumulative interface area build-up of individual interfaces during translation. The midpoints (dots) are residues at which half of the eventually formed area is exposed. Assembly-onsets are mapped to the nearest midpoint on condition that it is not a homomeric interface. (**B**) Pairwise comparison of cotranslationally forming (in the simultaneous mode) interfaces of multi-interface heteromeric subunits to the mean of all other heteromeric interfaces on them. For visual aid, see line diagram under panel C. Error bars represent SEM and labels on bars show the number of proteins in each group. The *p*-value was calculated with a Wilcoxon signed-rank test. (**C**) Pairwise comparison of the normalised assembly order in 201 complexes between cotranslationally forming and all other heteromeric interfaces. The normalised assembly order is a 0-to-1 scale where 0 and 1 represent the first and the last step of the predicted assembly pathway. The *p*-value was calculated with a Wilcoxon signed-rank test. (**D**) Two examples of simultaneous cotranslational assembly between subunit pairs in heteromeric complexes: the subunits E and B1 of the V-type ATPase (pdb: 6wm2), and importin-9 with histone H2A (6n1z).

Using this method, we identified 281 interfaces on multi-interface heteromeric subunits that may form in a simultaneous cotranslational fashion. To see if these correspond to the largest interfaces within each protein, we calculated the mean interface area for all other interfaces on these subunits to be able to perform a paired statistical test. Our results show that the identified cotranslationally assembling interfaces are indeed larger by 19% than other interfaces on these subunits (**Figure 2B**; *p* = 0.018, Wilcoxon signed-rank test).

We wished to put these interfaces into the context of their full complexes. Do simultaneously forming interfaces represent early forming subcomplexes that then initiate further assembly events, since the first step of a protein complex assembly pathway is the most likely to occur cotranslationally (Wells, Bergendahl and Marsh, 2015)? Although the largest interface in a complex is always predicted to assemble earliest in the assembly pathway, subsequent steps are non-trivial because they can involve multiple subunit:subunit interfaces (Ahnert et al., 2015; Levy et al., 2008; Marsh et al., 2013). To answer this question, we predicted the assembly steps of heteromeric complexes on the basis of their structures (Wells, Bergendahl and Marsh, 2016). This analysis revealed that the identified interfaces tend to form much earlier than other heteromeric interfaces in the complexes (**Figure 2C**; *p* = 0.01, Wilcoxon signed-rank test). Another interpretation of this can be given by classifying assembly steps into “early” and “late”, depending on their normalised assembly order (McShane *et al*., 2016), which is a 0-to-1 scale indicating the first-to-last steps of a pathway, where we defined early steps with values less than or equal to 0.5. According to this, a simultaneously forming interface is 1.7-times more likely to form early (180 [67%] of 270 *vs* 633 [54%] of 1171; *p* = 9.5 × 10^-5^, Fisher’s exact test).

Some of the identified interfaces belong to complexes that have been shown to use cotranslational assembly routes, such as the proteasome (Panasenko *et al*., 2019) and subunits of the transcription initiation complex (Kamenova *et al*., 2019). However, many are not yet described in the literature, for example, the loading of histone H2A onto importin-9 (**Figure 2D**) (Padavannil *et al*., 2019), which has been reported to act as a storage chaperone while transporting a histone dimer to the nucleus. Another example is the V-type ATPase (**Figure 2D**), whose catalytic A and B subunits have been tested for their ability to assemble in the sequential mode with a negative result (Shiber *et al*., 2018), but our structural approach using the assembly-onset identified the E1 subunit to form in the simultaneous mode with the catalytic B subunit.

### N-terminal interfaces tend to be larger than C-terminal interfaces supporting evolutionary selection for cotranslational assembly

There are two possible explanations for the observation that cotranslationally forming interfaces tend to be larger. First, larger interfaces may be inherently more likely to form cotranslationally because their assembly is more energetically favourable. In this scenario, cotranslational assembly has not been evolutionarily selected for; instead, the larger interfaces are simply more likely to be formed while the protein is still in the process of being translated, without providing any functional benefit. Alternatively, cotranslational assembly may have been selected for, e.g. because it increases the efficiency of assembly and avoids potentially damaging non-specific interactions. Here, large interfaces have evolved to increase the level of functionally beneficial cotranslational assembly.

One way to distinguish between these two scenarios is to compare the sizes of N and C terminal interfaces. Regardless of whether cotranslational assembly occurs simultaneously (**Figure 1B**) or sequentially (**Figure 3A**), due to vectorial synthesis on the ribosome, N-terminal regions of proteins are more likely to be involved in binding events during translation. Therefore, if cotranslational assembly is adaptive, we would expect that N-terminal interfaces in multi-interface heteromeric subunits, which will be translated first, should show a significant tendency to be larger than C-terminal interfaces, as illustrated in **Figure 3B**.

**Figure 3.**
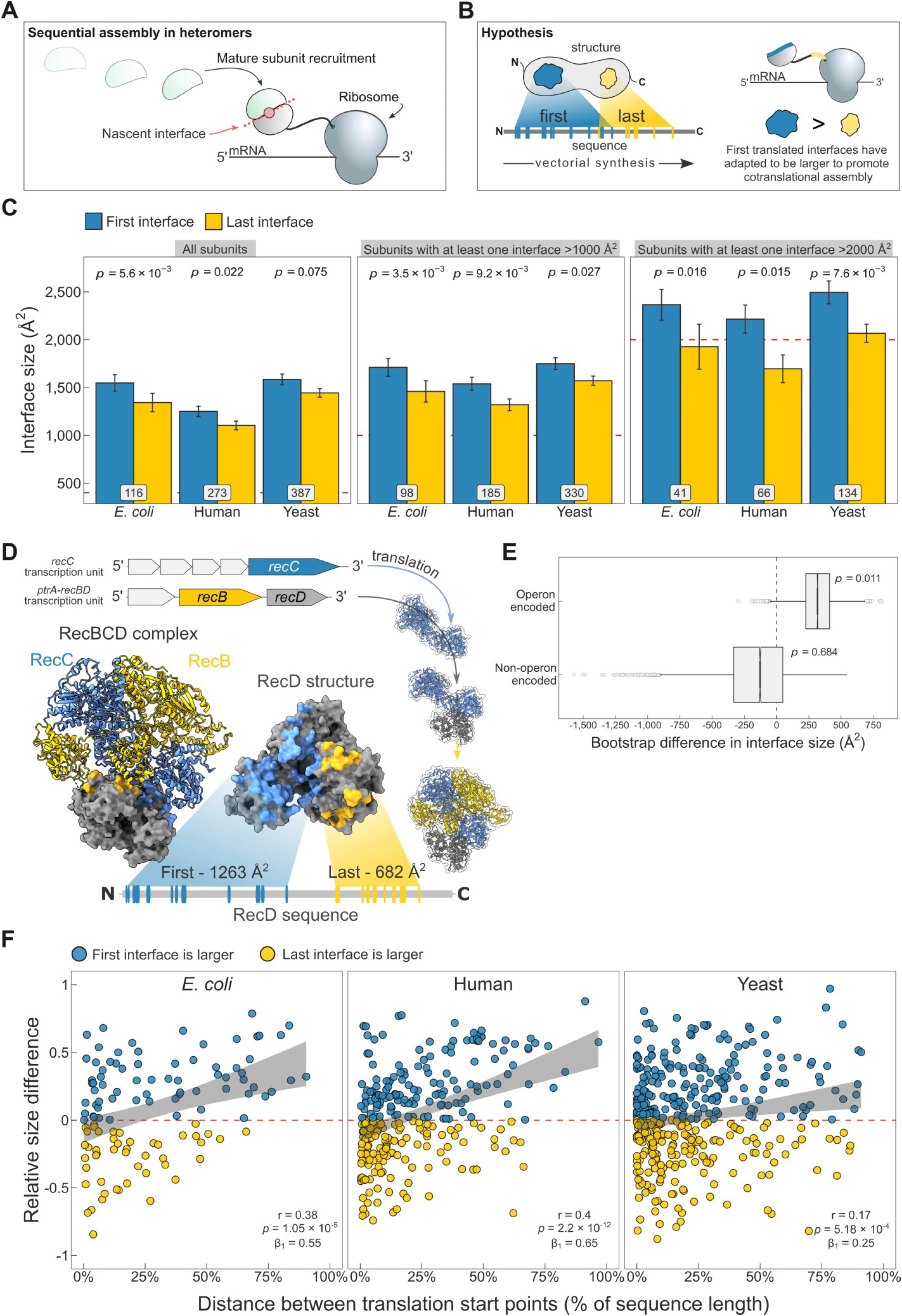
(**A**) Schematic representation of sequential cotranslational assembly in homomers. (**B**) Diagrammatic representation of the hypothesis test of the adaptive model of cotranslational assembly. (**C**) Area differences between the first and last translated interfaces in multi-interface heteromeric subunits across the species. Panels are ordered by the area cutoffs, 400, 1000, and 2000 Å^2^, which are satisfied if either the first or the last interface is larger than the cutoff. Error bars represent SEM and labels on bars show the number of proteins in each group. The *p*-values were calculated with Wilcoxon signed-rank tests. (**D**) Example of an operon-encoded complex, the RecBCD nuclease (pdb: 5ld2). In the linear sequence of RecD, the interface with RecC is translated first, and that with RecB is last. The RecD:RecC interface is twice the area of the RecD:RecB interface, likely to promote cotranslational subunit recruitment. (**E**) Bootstrap distributions of the area difference between the first and the last translated interface in operon and non-operon encoded complexes. The *p*-values were calculated from 10^4^ resamples with correction for finite testing. (**F**) Correlation between the relative distance of translational start points and the relative area difference of the first and last translated interfaces. Shaded lines represent the 95% confidence interval of the regression line. The Pearson’s correlation coefficient *r*, its *p*-value, and the regression coefficient β1 are shown in the panels.

To address the problem, we selected experimentally determined heteromeric complex structures from three model proteomes: *Escherichia coli (E. coli), Saccharomyces cerevisiae* (yeast), and *Homo sapiens* (human). We also supplemented the smaller yeast data set with heteromeric models computed for yeast core complexes, which were determined using residue coevolution inferred from paired multiple sequence alignments, followed by deep learning-based structure prediction of subunit pairs with experimental evidence to interact (Humphreys *et al*., 2021). For each heteromeric subunit, we defined the first interface as the one that exposes the first interface residue in the linear protein sequence, and, to treat the termini symmetrically, the last interface was defined as the one that exposes the last interface residue, *i.e*. the first interface residue from the C-terminal direction.

When we compare the areas of the first and last translated interfaces in heteromeric subunits across species, we find the first interface to be larger (**Figure 3C**; full distribution in **Figure S2A**). The strongest effect is measured in *E. coli*, where the first interface is 15% larger on average than the last (*p* = 5.6 × 10^-3^, Wilcoxon signed-rank test). In human multi-interface heteromeric subunits, the difference is 13% (*p* = 0.022). A weak trend is also detected in the yeast data set, where the first translated interface is overall 10% larger. However, since larger interfaces tend to cotranslationally assemble more frequently, we filtered the subunits to include cases where the size of at least one of the two interfaces is larger than 1000 or 2000 Å^2^, with the assumption that subunits containing only small interfaces would be unlikely to undergo any cotranslational assembly (**Figure 3C**). This analysis revealed a strong proclivity of the first translated interface to be larger than the last when the probability of cotranslational assembly is higher. The mean differences are 15%, 17%, and 23% in *E. coli*, 13%, 17%, and 31% in human, and 10%, 11% and 21% in yeast, respectively to the increasing area thresholds.

Our attention was next drawn to *E. coli*, which demonstrated the strongest trend. In prokaryotes, many heteromeric complexes are encoded by operons, where the different subunits are translated off of the same polycistronic mRNA molecules. Early studies in bacteria indicated that operon gene order is correlated to physical interactions between the encoded proteins (Mushegian and Koonin, 1996; Dandekar *et al*., 1998). Further investigation laid down theoretical, mechanistic, and evolutionary evidence in support of this (Sneppen *et al*., 2010; Shieh *et al*., 2015; Wells, Bergendahl and Marsh, 2016). Cotranslational assembly is likely to be particularly common in operon-encoded heteromers, given that the translation of different subunits is inherently colocalised. We hypothesised that the tendency for N-terminal interfaces to be larger should be stronger in operon-encoded *E. coli* heteromers, compared to those that are not operon encoded heteromers.

We illustrate the example of the RecBCD nuclease in **Figure 3D**. Genes of the subunits are located in adjacent loci encoding transcriptional units for RecC and RecB/D. One study reported that purification of RecD is complicated by the formation of inclusion bodies, while the other two subunits remain in the soluble fraction (Masterson *et al*., 1992). Moreover, a genetic analysis proposed that partially folded RecC and RecD might interact during translation, or that RecC forms a complex with RecB first, onto which RecD is then assembled (Amundsen, Taylor and Smith, 2002). The regulatory subunit RecD has two interfaces well separated in the sequence, where the interface with RecC is translated first. One might imagine that the nascent chain of RecD forms a complex with mature subunit of RecC, having double the interface area to accommodate RecC than that for RecB. In this scenario, the assembly efficiency is not only maximised by gene order reducing stochasticity, but also by cotranslational assembly minimising the need for post-translational association.

To further test whether large interfaces could have been selected to promote cotranslational assembly, we acquired annotations derived from RNA sequencing datasets (Chetal and Janga, 2015) to group heteromers from *E. coli* according to whether or not they are encoded by operons. We generated bootstrap distributions of the area difference between the first and the last translated interface to visualise and derive a probability (**Figure 3E**). In agreement with the above, we found that the area difference between the two interfaces is significantly larger in operon-encoded multi-interface heteromeric subunits, favouring the first interface (mean area difference of 317 Å, *p* = 0.011).

We speculate that the first *versus* last interface trend may be the hallmark of sequential cotranslational assembly (**Figure 3A**), rather than that of the simultaneous mode (**Figure 1B**). The strong trend in *E. coli* supports this idea, because polycistronic gene structure is more compatible with sequential assembly. In eukaryotes, large complexes and subunits of lowly abundant complexes may require an additional biological process to ensure their transcripts are colocalised for simultaneous assembly and to facilitate further assembly steps (Keene, 2007; Chen and Mayr, 2022). Sequential assembly, on the other hand, may have evolved to exploit large interface areas for the recruitment of partner subunits. This can be conceptualised as the “bait and prey strategy” of cotranslational assembly, in which a large nascent interface represents the “bait” that is then bound by a fully folded “prey” subunit. To substantiate this model, we removed from the human dataset those proteins that were identified in the previous section as having simultaneously forming interfaces. Strikingly, removal of these proteins increases the difference between the first and the last translated interface from 13% to 18% (**Figure S2B**; *p* = 0.014, Wilcoxon signed-rank test). One explanation is that these heteromers are enriched in sequential assembly, and thus exhibit a greater difference.

A property that would be consistent with the above model is interface separation. The later the translation of the last interface starts relative to the first, the higher the chance that assembly of the first interface will be undisturbed, free of competition with the partner subunit of the last interface. Therefore, we hypothesised that the distance between translation start points of the first and last interface, which are the earliest emerged interface residues of each, should correlate with the size difference in favour of the first interface. Because of large variances in protein length and interface size, we normalised the translational distance between the first and last interface as the percentage of the protein’s sequence length, and scaled the area difference by the sum of both interfaces. **Figure 3F** shows the correlation between the separation of translation start points and the area differences of the first and the last interface (absolute values in **Figure S2C**). As expected, increasing the distance between the start points monotonically increases the extent of the area difference across all species. In case the interface separation metric is confounded by sequence length, we split the structures into less and more than 400 amino acids, which is the pan-species mean of sequence lengths. In both subsets, there is a pronounced preference in all species for a larger first interface when the separation is high (**Figure S2D**). The causal direction of this effect, whether it reflects that cotranslational assembly happens more often in high degrees of interface separation, or that separation is driven by selection for cotranslational assembly, remains to be answered.

## Discussion

It has long been understood that interface area is important for assembly, but the capacity in which it shapes the hierarchy of individual interfaces on subunits remained elusive. In this study, we first combined information from ribosome profiling with structural data on protein complexes to probe the importance of interface area in the process of cotranslational assembly. Our results demonstrate a strong correspondence between interface size and cotranslational subunit binding. Inspired by this, we set out to test an important question about the biological significance of cotranslational assembly: do large interfaces give rise to cotranslational assembly because of simple energetic reasons or do they reflect an evolutionary adaptation for a functional benefit? We found a clear trend across three evolutionarily distant species for the first translated interface of heteromeric subunits to be larger, suggesting that large interfaces have evolved to promote cotranslational assembly.

While our results support the adaptive hypothesis of interface size, it is not entirely inconceivable that cotranslational assembly represents a ratchet-like mechanism of constructive neutral evolution (Gray *et al*., 2010; Hochberg *et al*., 2020), whereby a drift in interface properties creates ideal conditions for assembly on the ribosome. Reversion to the post-translational route is prevented by the accumulation of mutations that are neutral in the subunit entrenched in cotranslational assembly, but would otherwise be deleterious in the ancestor. A similar neutral process may sustain the differences in interface area presented in this study.

In light of current knowledge of bacterial operon structure and translation regulation, it is not surprising that we observed the strongest trend among *E. coli* heteromers with respect to the size of the interface that first emerges from the ribosome. Supposedly, the effect is attributable to the widespread sequential assembly between mature subunits and nascent chains, reflecting the mechanism of effective subunit recruitment by large N-terminal “bait” interfaces. An interesting question for laboratory experiments is whether operon-encoded heteromers can assemble in the simultaneous mode, providing that the structural organisation of the bacterial polysome allows for such a precise coordination (Brandt *et al*., 2009)?

How does a large interface area help translating ribosomes find one another? Its benefit in homomers for facilitating cotranslational assembly is clear, because the subunits are localised to the same mRNA, and large interfaces are more likely to form interactions before translation is complete. In heteromers, one hypothesis argues the involvement of RNA binding proteins that orchestrate transcript colocalisation (Keene, 2007; Chen and Mayr, 2022), from where similar rules may apply to heteromer assembly as for homomers. A more parsimonious hypothesis is formed on the observation that cotranslational assembly can result in transcript colocalisation, which is ablated when subunit affinity is decreased (Heidenreich *et al*., 2020). This may suggest that affinity, which correlates strongly though imperfectly with interface size (Brooijmans, Sharp and Kuntz, 2002; Vangone and Bonvin, 2015), can play a role in the colocalisation of transcripts belonging to the same complex.

Many more topics of inquiry remain open for future studies. Analogous to protein folding, cotranslational assembly can be thought of as a hydrophobic collapse that shapes the quaternary structure of the complex. As with folding in the cell, the involvement of other factors must not be overlooked. A wide array of ribosome-associated chaperones are vital for nascent chain homeostasis (Oh *et al*., 2011; Döring *et al*., 2017; Koldewey, Horowitz and Bardwell, 2017; Shiber *et al*., 2018; Zhang *et al*., 2020) and the degree to which they choreograph assembly steps is yet to be elucidated.

Attention should be paid to the far-reaching genetic consequences of cotranslational assembly (Natan *et al*., 2017). How much does transcript-specific assembly buffer the dominant-negative effect, and what does it mean in the context of human genetic disease (Bergendahl *et al*., 2019)? This effect requires mutant subunits to be stable enough to assemble into complexes, and thus the impact of the mutations tends to be milder at the structural level than of other pathogenic mutations (McEntagart *et al*., 2016), making them exceptionally difficult to detect using the existing variant effect predictors (Gerasimavicius, Livesey and Marsh, 2021). Interestingly, cotranslational assembly should actually make the dominant-negative effect less common in homomers, because it can limit the mixing that occurs between wild-type and mutant subunits. It remains to be seen whether dominant-negative mutations are in fact less common in cotranslationally assembling complexes.

Finally, our results build on evidence from the past decade and emphasise the importance of protein complex assembly at the translatome level. Although evolutionary selection against N-terminal interface contacts to avoid premature assembly was previously found in homomers (Natan *et al*., 2018), here we report an opposite phenomenon in which proteins that do cotranslationally assemble sustain large N-terminal interfaces in order to promote cotranslational subunit recruitment.

## Methods

### Protein structural datasets

Starting from the entire set of structures in the Protein Data Bank (Berman et al., 2000) on 2021-02-18, we searched for all polypeptide chains longer than 50 residues with greater than 90% sequence identity to *Homo sapiens, Saccharomyces cerevisiae*, and *Escherichia coli* canonical protein sequences. When proteins mapped to multiple chains, we selected a single chain sorting by sequence identity, then number of unique subunits in the complex, and then the number of atoms present in the chain. Only biological assemblies (pdb1) were used and symmetry assignments were taken directly from the PDB. Polypeptides formed by cleavage were excluded. In the generation of the multi-interface heteromeric subunit data sets, chains with an at least 70% complete structure were considered and only included if they formed interface pairs >800 Å^2^ with at least two different subunits. The computed structures of yeast core complexes were downloaded from the ModelArchive link provided by Humphreys et al., 2021. For downstream analysis, mmCIF files were converted into pdb format and the chains were mapped to genes using the table provided on ModelArchive. In the analysis of N- vs C-terminal interface sizes, we excluded very large complexes, defined as those containing ≥10 subunits. This is because of the previous evidence that predicting assembly order based on interface size in very large complexes is not as accurate (Marsh *et al*., 2013; Ahnert *et al*., 2015), likely because of the many intersubunit interfaces these complexes possess.

### Computation of interface related properties

Pairwise interfaces were calculated at residue-level between all pairs of subunits using AREAIMOL from the CCP4 suite (Winn *et al*., 2011) with the default surface probe radius of 1.4 Å. The interface was defined as the difference between the solvent accessible surface area of each subunit in isolation and within the context of the full complex. An interface area cutoff of >400 Å^2^ was used for homomeric subunits and individual interfaces of multi-interface heteromeric subunits derived from the PDB to exclude potential crystallographic interfaces and to restrict the analyses to biologically significant interfaces. Assembly order was computed by predicting the assembly pathway assuming additivity of pairwise interfaces in each complex (Marsh *et al*., 2013), and implemented with the *assembly-prediction* Perl package (Wells, Bergendahl and Marsh, 2016).

The relative interface location was calculated according to the formula:

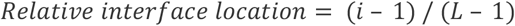

where *i* marks the residue at which half of the cumulative buried surface area of the interface is passed (interface midpoint), and *L* is the sequence length.

The normalised distance between translational start points was calculated as:

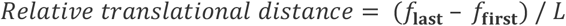

where *f* marks the first residue of the given interface and *L* is the sequence length.

Area differences between the first and the last interface were normalised according to the equation:

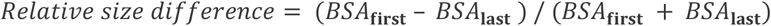

where *BSA* is the buried surface area of the corresponding interface.

### Mapping simultaneously forming interfaces in multi-interface heteromeric subunits

Cotranslational assembly-onset positions were acquired from the supplemental material of Bertolini et al., 2021. From the onset positions, 30 residues were subtracted to account for the length of the ribosome tunnel. Our method mapped the assembly-onset position to the closest interface midpoint in the linear sequence. Cases where the assembly-onset mapped to a homomeric interface were discarded under the assumption that the homomeric interface is hierarchically higher and undergoes *cis*-assembly. In subsequent analyses presented in **Figure 2B/C**, we only included comparisons between heteromeric interfaces.

### Plasma membrane localisation

We obtained plasma membrane annotations directly from the UniProt FTP site (Bateman *et al*., 2021). UniProt entries with the gene ontology term “GO:0005886” (plasma membrane) were filtered for canonical sequences.

### Mapping subunits to operons

Operon annotations were downloaded from OperomeDB (Chetal and Janga, 2015). Genes were mapped to UniProt identifiers using *E.coli* proteome-specific mapping from the UniProt FTP site (Bateman *et al*., 2021).

### Molecular graphics

Visualisation of structures was performed with UCSF ChimeraX version 1.1 (Pettersen *et al*., 2021).

## Statistical analysis

Data exploration and statistical analyses were carried out in RStudio (Rstudio, 2020) version 2022.02.0+443 “Prairie Trillium” Release, using R version 4.1.2 (R Core Team, 2014). The R packages used for analyses are `tidyverse`, `tidytable`, `rsample`, `rstatix`, `scales`, and `ggbeeswarm`. The Wilcoxon rank-sum or signed-rank tests were used for A/B testing of interface size distributions, because although they appear log-normal, they are also left-bounded due to the minimum interface area filter, thus a non-parametric test was required. Wilcoxon signed-rank tests were one-tailed, and their main assumption that the data are symmetric around the median was supported by boxplot distributions. In the bootstrap analysis, the data was stratified for operonal localisation in 10^4^ resamples. The *p*-value was calculated by determining the fraction of point estimates (difference in area) greater than 0, with correction for finite sampling. In the regression analysis, conditions for statistical inference, including linearity of the relationship between variables, the independence and normality of the residuals, and homoscedasticity were met; validations can be found in the analysis script.

## Data and code availability

Data and code to reproduce the results have been deposited on the OSF at https://osf.io/x5b2n/.

## Acknowledgements

MB is supported by the Biotechnology and Biological Sciences Research Council EASTBIO DTP (BB/M010996/1), and used resources provided by the Edinburgh Compute and Data Facility. JAM was supported by a Medical Research Council Career Development Award (MR/M02122X/1) and is a Lister Institute Research Prize Fellow.

## Author contributions

JAM conceived the project and generated the structural and assembly order data. MB acquired all other data for use in the study and carried out the analysis. JAM supervised and aided in interpreting the results. MB wrote the manuscript, which was finalised with considerable contributions by JAM.

---

**Figure S1.**
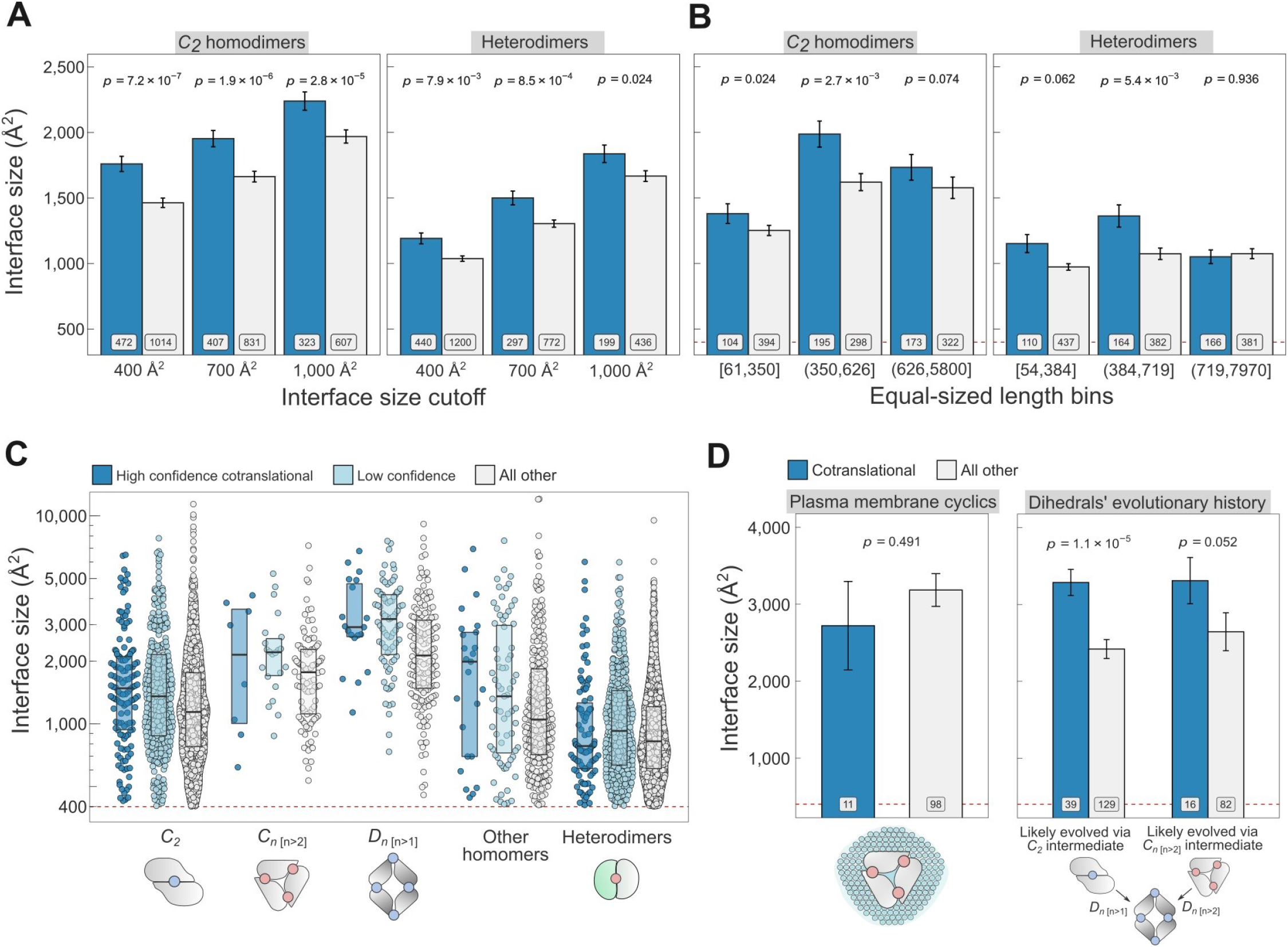
(**A**) Interface size differences between cotranslationally assembling and all other subunits of *C_2_* homodimers and heterodimers, measured at incremental interface area cutoffs. Error bars represent SEM and labels show the number of proteins in each group. The *p*-values were calculated with Wilcoxon rank-sum tests. (**B**) Interface size differences between cotranslationally assembling and all other subunits of *C_2_* homodimers and heterodimers, binned into 3 approximately equal-sized bins of sequence length. Error bars represent SEM and labels show the number of proteins in each group. The *p*-values were calculated with Wilcoxon rank-sum tests. (**C**) Interface size differences subset by confidence in cotranslational assembly. Pictograms show the basic structure of symmetry group members, with the blue dots representing isologous and red dots heterologous and heteromeric interfaces. (**D**) *Left:* Interface size differences between cotranslationally assembling and all other subunits of plasma membrane localised higher-order cyclic symmetry members. *Right:* Interface size differences between cotranslationally assembling and all other subunits of dihedral complexes grouped by their probable evolutionary history. The *p*-values were calculated with Wilcoxon rank-sum tests.

**Figure S2.**
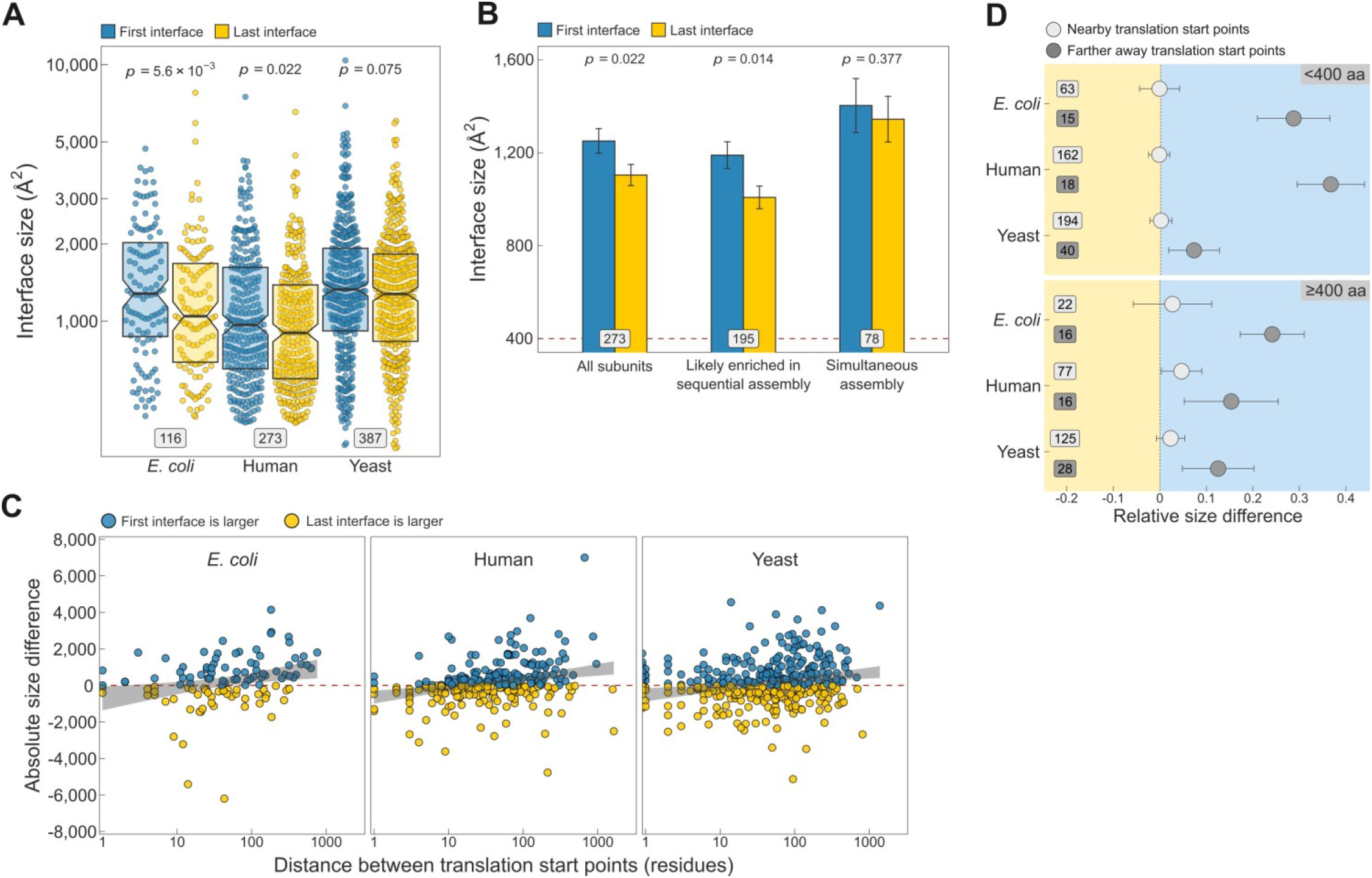
(**A**) Full distribution of the first *versus* last translated interface size trend shown in the first panel of **Figure 3C**. (**B**) Interface size differences between the first and last translated interfaces in human multi-interface heteromeric subunits, with those identified to have simultaneously forming interfaces shown as a separate group. Error bars represent SEM and labels on bars show the number of proteins in each group. The *p*-values were calculated with Wilcoxon signed-rank tests. (**C**) Scatter plots showing the absolute distance in amino acids between translational start points of the first and last translated interfaces and the absolute area difference across species. Shaded lines represent the 95% confidence interval of the regression line. (**D**) TIE-fighter plots demonstrating that interface separation increases the area difference in favour of the first interface independent of protein length. For all species, the relative translational distance interval was split at the mean, and the plot is divided into less and more than 400 amino acid long sequences. Dots represent the mean and error bars are SEM. Labels are the number of proteins in each group. Background colour reflects the direction of the size difference: blue – first interface larger, yellow – last interface larger.

